# Automatic Detection of Larval Zebrafish ECG: Computational Tool for High-throughput Cardiac Activity Analysis

**DOI:** 10.1101/2021.02.08.430220

**Authors:** Richard Barrett, Rhiannon Hurst, Edward Tarte, Ferenc Müller, Attila Sik

## Abstract

Automatic analysis of larval zebrafish electrocardiographs (ECG) is essential for high-throughput measurements in environmental toxicity assays, cardiotoxicity measurements and drug screening. We have developed a MATLAB based software is built on methods that have previously been used to analyze human ECG, such as the Pan-Tompkins algorithm and Wavelet. For the first time these sophisticated tools have been applied to larval zebrafish ECG to automatically characterize the heart-beat waveforms. The ability of the automated algorithm to detect the QT interval for both normal and pharmacologically altered larval ECG is found and compared to previously used software that is built into LabChart^®^ (AD Instruments). Using wavelet transforms it is shown that the typical larval ECG features are within the frequency range of 1 to 31 Hz. It is also shown that the automated software is capable of detecting QTc (heartrate corrected heartbeat interval) changes within pharmacologically altered zebrafish larval ECG. The automated process is a significant improvement on the approaches that were previously applied to the zebrafish ECG. The automated process described here that is based on established techniques of analyzing ECG can sensitively measure pharmacologically induced changes in the ECG. The novel, automated software is faster, more sensitive at detecting ECG changes and less dependent on user involvement, thus minimizing user error and human bias. The automated process can also be applied to human ECG.

## Introduction

Using zebrafish larvae for chemical compound screening is becoming increasingly important for cardiac drug development. It has been previously shown that the zebrafish electrocardiogram (ECG) is similar to mammals. Another advantage of the zebrafish model is that a minimal amount of chemicals are necessary for drug testing and embryo production is fast and inexpensive [1]. Furthermore, using mammals for preliminary screening is expensive, slow and requires enormous numbers of animals. Video-recording of zebrafish embryos’ heart activity is currently used for drug screening, but unlike ECG recording it lacks the temporal and dynamic resolution necessary for cardiac cycle component analysis [2–4]. To aid development of high-throughput, high fidelity methods to simultaneously and automatically record ECGs from larval zebrafish, analysis tools that automatically characterize larval ECG signals under control conditions and after drug treatment are desirable.

Presented herein is the first successful attempt to develop analysis tools based on algorithms and methods that have previously been used to characterize human ECG, including the Pan-Tompkins algorithm and wavelet transform analysis implemented in MATLAB. These tools are compared to techniques that have previously been used to analyze zebrafish larval ECG based on the software packages offered by AD Instruments (LabChart^®^).

## Methods

### Data acquisition

The data used to investigate the analysis tools is larval zebrafish ECG that was taken from 3 days post fertilization larvae using the methods outlined in [5]. The baseline ECG was recorded for approximately 2 minutes before verapamil was added and the ECG was recorded for a further 25 minutes. This data represents the typical ECG of a zebrafish larvae recorded in our lab and a typical drug response as previously demonstrated in our previous paper [5].

### Data Extraction

After the recording, the data was split into 8, 1-minute sections, labelled chronologically as A, B, C, D, E, F, G, H and exported from LabChart^®^ as a MATLAB compatible data format. Section A was taken from ECG recorded before the drug was introduced to the solution whereas sections B-H were recorded afterwards. Each section was analyzed separately using the different tools outlined above to determine parameters such as the heart rate and the QTc.

### Down-sampling the Data for MATLAB Programs

All of the programs written for MATLAB use some initial down-sampling to speed up the processing of the algorithms. Assuming that all the relevant information in the zebrafish ECG occurs well below 100 Hz the program down samples to a sampling rate of *f_s_*=200 Hz. In the MATLAB code this appears as:

%MATLAB ALGORITHM TO DOWNSAMPLE ECG
%% Down-sample to 200 Hz sampling rate
out =~rem(FS,200)*FS/200;

### Determining the frequency of the components of the zebrafish ECG

To investigate which filtering bandwidth would best capture the ECG features whilst removing any low-frequency drift and high-frequency interference, the sections were first analyzed using a wavelet transform in MATLAB. To perform the transform, 20 seconds of raw ECG data was taken from section A and transformed using a Gaussian wavelet combined with an in built MATLAB function, as described in above. The result of the wavelet transform was outputted as a contour map and an approximate of the characteristic frequencies of the ECG features were measured visually. Due to space-time localization it is not possible to state categorically that the characteristic frequency of each feature was perfectly aligned to the correct time signature, however the graphical output could be used as a guide of the approximate frequency of each feature. This frequency of each R wave and T wave within the section was recorded and tabulated. Graphical outputs of the ECG wavelet transform. From the tabulated data the mean and standard deviation of the characteristic frequency of the R and T waves were determined. These values were used to tailor the upper frequency cut-off to remove higher frequency noise without attenuating the ECG signal. The upper frequency limit was set to the mean R-wave characteristic frequency plus two standard deviations. Assuming that the R-wave frequency is normally distributed about the mean, this would suggest that approximately 95% of the R-waves would be captured without attenuation. The lower frequency cut-off was set to 1 Hz for reasons that are outlined above.

### LabChart^®^ Analysis of Sections

To analyze the data through the inbuilt ECG analysis in LabChart^®^, each section was first band-pass filtered using the upper and lower cut-offs. The filtered ECG was then analyzed using the protocol outlined and Fig 1.

**Fig 1.**
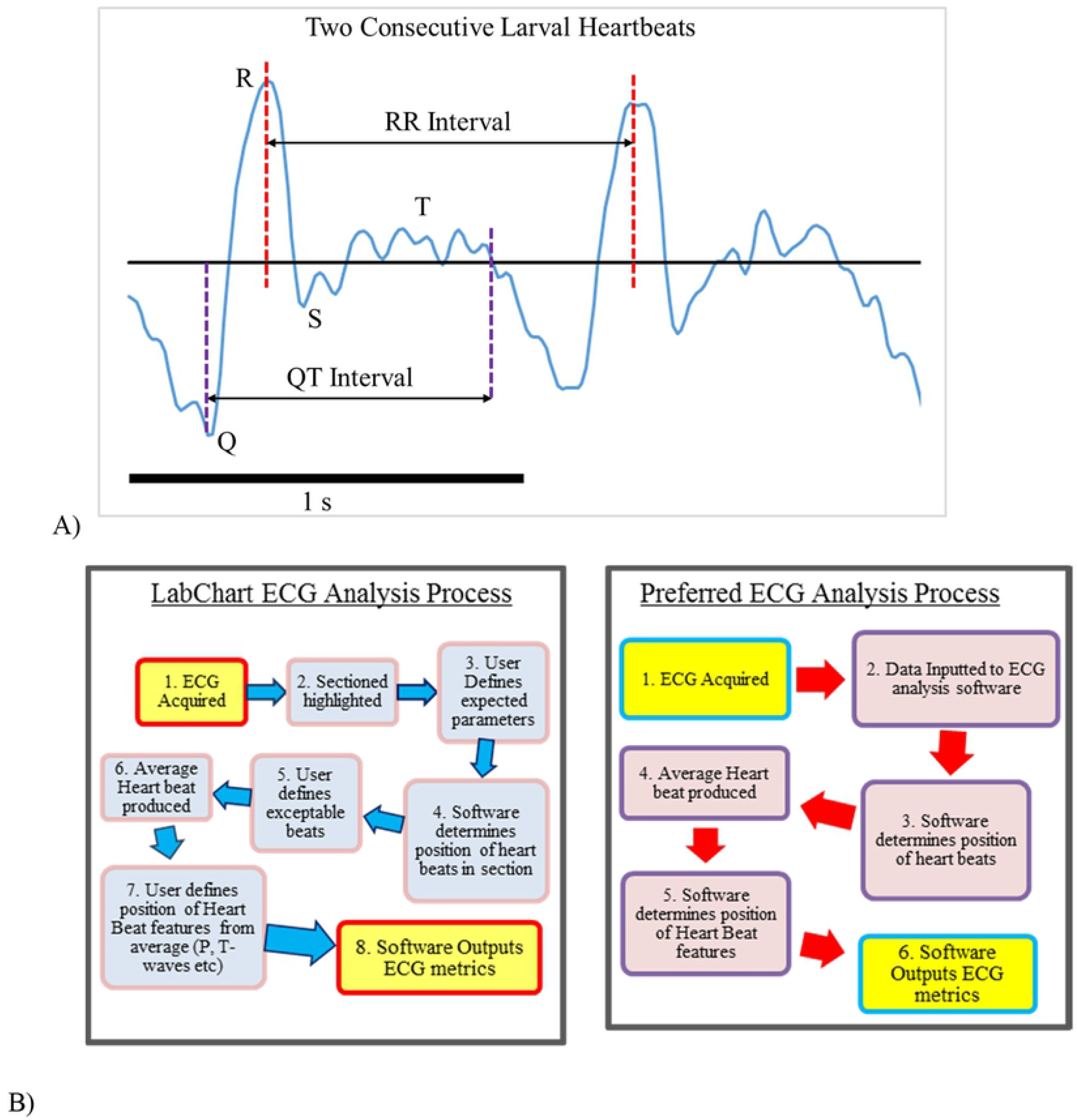
Zebrafish larval heartbeats and strategies of the analytical methods. A) Two consecutive larval heart beats that have been annotated to show the position of the Q, R S and T-waves. B) The two strategies employed to analyse the ECG of Larvae. Left: The standard analysis protocol that is part of the inbuilt LabChart^®^ analysis software. Right: The automated analysis protocol that has been implemented in MATAB

The LabChart^®^ software automatically locates the position of each R-wave based on user inputs such as, typical QRS-width, typical RR-interval and QT-interval. Thus the analysis requires some user involvement from the start. After the location has of each heartbeat has been found it is then decided by the user which heartbeats should be deemed ‘acceptable’. This is included in the software so that anomalous or corrupted recording can be omitted from the overall average. In the larval ECG that was studied there were no anomalous beats and so each beat was accepted. This information is then used by the software to produce and average waveform that is produced by finding the mean voltage of the heartbeat at each time point, relative the position of the R-wave.

From this average view, the user defines the position of the Q-start point, Q-end point, T-peak and T-end. From this input the software automatically calculates the QTc (based on the Bazett formula) and other heartbeat characteristic. This information was exported as a table into Microsoft Excel for further analysis. Performing the analysis on a section of data takes at least 5 minutes.

This process was performed on each section of larval ECG to determine the heartbeat characteristics before and after the introduction of the pharmacological agent.

#### Applying the Automated process

The automated process was applied to each data section and the outputted QTc was then exported to Excel. The program measured the QTc for each 40 consecutive heartbeats and the output was tabulated as shown in Table I.

**Table 1.**
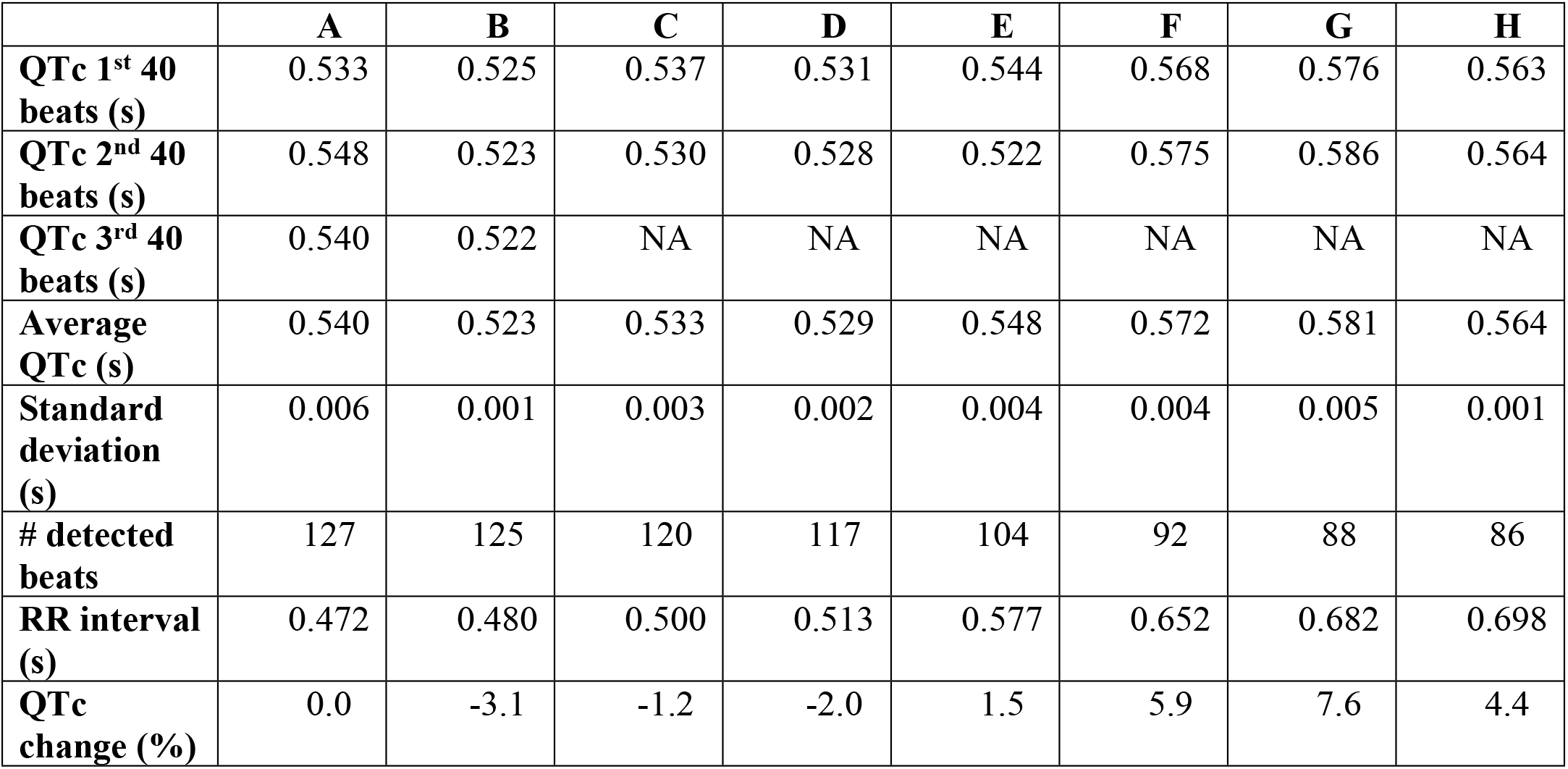
The outputted data from automated process that has been collated

|  | **A** | **B** | **C** | **D** | **E** | **F** | **G** | **H** |
| --- | --- | --- | --- | --- | --- | --- | --- | --- |
| **QTc 1<sup>st</sup> 40 beats (s)** | 0.533 | 0.525 | 0.537 | 0.531 | 0.544 | 0.568 | 0.576 | 0.563 |
| **QTc 2<sup>nd</sup> 40 beats (s)** | 0.548 | 0.523 | 0.530 | 0.528 | 0.522 | 0.575 | 0.586 | 0.564 |
| **QTc 3<sup>rd</sup> 40 beats (s)** | 0.540 | 0.522 | NA | NA | NA | NA | NA | NA |
| **Average QTc (s)** | 0.540 | 0.523 | 0.533 | 0.529 | 0.548 | 0.572 | 0.581 | 0.564 |
| **Standard deviation (s)** | 0.006 | 0.001 | 0.003 | 0.002 | 0.004 | 0.004 | 0.005 | 0.001 |
| **# detected beats** | 127 | 125 | 120 | 117 | 104 | 92 | 88 | 86 |
| **RR interval (s)** | 0.472 | 0.480 | 0.500 | 0.513 | 0.577 | 0.652 | 0.682 | 0.698 |
| **QTc change (%)** | 0.0 | -3.1 | -1.2 | -2.0 | 1.5 | 5.9 | 7.6 | 4.4 |

The program also generated a plot of the average waveform found for each set of 40 beats, and searched for Q-start, Q-end, T-peak and T-end. These intervals were fed into the Bazett formula to calculate the QTc for each set of beats. An overall QTc for each section of data was found by taking the mean of the measured QTc for each set of beats. The time taken to analyze each section was approximately 10 seconds.

## Results

### Larval ECG

We have previously reported larval ECG capture for characterization of arrhythmia induced by pharmacological agents in zebrafish [5]. By acquiring ECG from these model organisms we are able to evaluate the cardiac effects of drug exposure, temperature changes, anesthetic exposure and staging. A typical larval ECG signal is shown in Fig 1A, above. Furthermore, the ECG offers a high fidelity output of the atrial-ventricular rhythmicity that is responsible for many arrhythmias in humans [6]. We have previously shown that this relationship can be altered in the zebrafish to mimic QTc prolongation in humans by adding cardio-active drugs such as verapamil [5]. However, to analyze the zebrafish ECG we have previously relied on software within the LabChart^®^ suite of programs for collecting electrophysiological data. Although the program is powerful, data analysis is a slow process and requires significant user involvement that potential introduces human error.

### LabChart^®^ Analysis Software

LabChart^®^ is a software package offered by AD Instruments to store and analyze electrophysiological measurements that are recorded using PowerLab hardware. The program has a number of inbuilt packages for analyzing ECG data and has previously been used to record and analyze larval zebrafish ECG. LabChart^®^ is used in this work to compare the analysis outputs versus the alternative tools under investigation. The LabChart^®^ ECG analysis package is heavily user-dependent as shown in Fig. 1B. The user must first select a period of ECG that they wish to analyze, specify the RR interval of the data and other characteristics. With this information the software is then able to determine the position of every R-wave within the ECG and attribute a fiducial mark to this time point. The user is then able to select or deselect detected “heart beats” based on their “Isoelectric Noise”, “Form Factor” or “RR-interval”, by setting an acceptable range and domain. Based on the user selections the software then averages all of the accepted heartbeats to produce an average waveform that represents the overall ECG activity for the selected time-period. From this averaged waveform the user determines the position of the P, Q and T features and the software is then able automatically calculate QTc (heartbeat interval corrected for heartrate) and other parameters of the ECG. By performing this process for larval zebrafish ECG before and after drug treatments, it is possible to evaluate any pharmacologically induced alterations in the heart beat cycle. However, this process is time consuming (approximately 3 minutes for 1 data section) and is heavily dependent on the users’ interpretation of the ECG signal.

### The Automated Analysis Software

#### Pan-Tompkins Algorithm for QRS Detection

To build automated signal processing software it is necessary to detect the QRS events of the larval ECG first. Most automated ECG programs attempt to detect the QRS first as it is a dominant feature that is the most robust to change in the cardiac cycle [7]. For this process a program has been developed based on the QRS detection algorithms established by Pan and Tompkins in the 1980’s which have been shown to be robust for many different types of ECG signals [8].

The original architecture of the Pan-Tompkins algorithm is divided into three processes which can be thought of as the Initial Learning Phase (ILP), Secondary Learning Phase (SLP), and Detection Phase (DP). The ILP initializes detection thresholds based upon the size of the “signal” and “noise” peaks detected. The SLP uses two full heart cycles to determine the average RR interval and then set the limit of the possible RR-interval for the rest of the ECG trace. To allow for any adaptation of the recorded ECG signal, for example due to pharmacologically induced change in heart rate, these thresholds are adjusted periodically. The DP processes the ECG and generates a pulse for each QRS. It then uses thresholds based on both the filtered ECG and the processed signal to detect the QRS waves. Their detection thresholds were set to just above the “noise” that is sensed by the algorithm. This approach reduced the number of false positives caused by noise that mimics the QRS characteristics, which is always a problem in human ECG due to electromyography artefacts. In zebrafish these artefacts could also be a problem due to sporadic motion artefacts caused by twitching or other motion. The automated process conserves and builds on this architecture.

Pan-Tompkins (P-T) used four simple algorithms to process the ECG data and one further algorithm to detect the QRS features from the processed data. These stages can be outlined as:

1. Filter the signal to remove artefacts, such as electrical noise at 50 Hz or baseline wander (< 1 Hz), and allow the signal to be processed efficiently. In their original work P-T used a transfer function to implement an Infinite Impulse Response Filter (IIR Filter) to remove the artefacts. Unfortunately, these types of filters introduce a phase lag that is proportional to the frequency of the input, and so is undesirable for this work. Instead, for our software inbuilt MATLAB functions are used to filter the signal without a phase lag. The MATLAB code used to implement the filter is shown below. As seen in the code, a filtfilt function is employed that applies a 3^rd^ order Butterworth filter to the raw data to band-pass the signal between the upper and lower frequency cut offs that are user defined. The filtered data is then normalized.

%MATLAB ALGORITHM TO FILTER RAW ECG
%Filt_low is user defined cut off to remove low frequency artefacts
%Filt_high is user defined cut off to remove high frequency artefacts
%ecg_raw is a vector that contains the sampled raw ECG recording
%FS is the sampling frequency
% ecg_filt is the filtered ECG signal

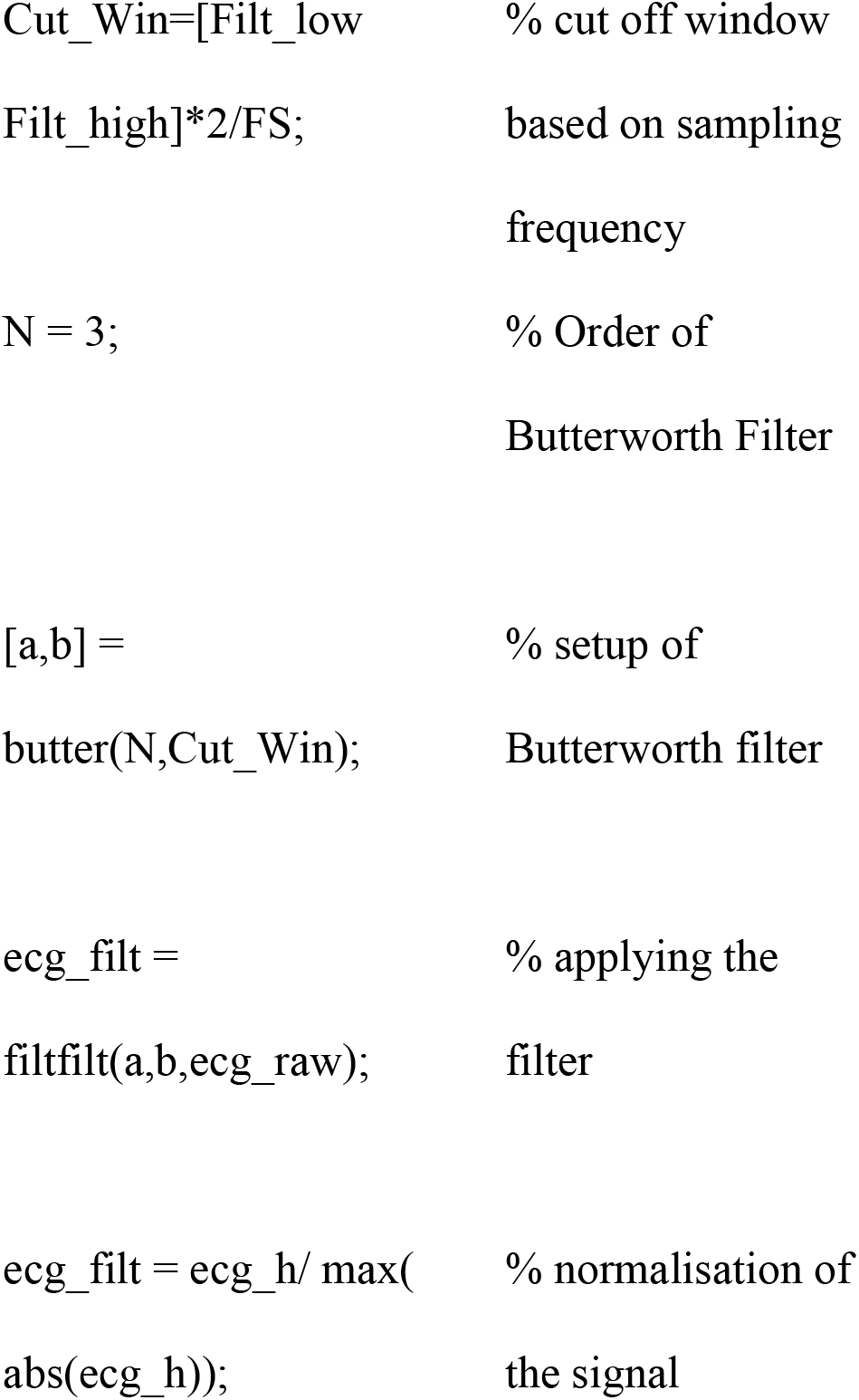
2. The second stage of the algorithm is to differentiate the signal to accentuate the turning points. If all artefacts have been removed then the turning points can only be associated with biological events, which for a normal ECG signal occur at the P-wave, R-wave and T-wave. By differentiating, these features become amplified. The amount that they are amplified is proportional to their frequency as higher frequency activity is changing at a faster rate.

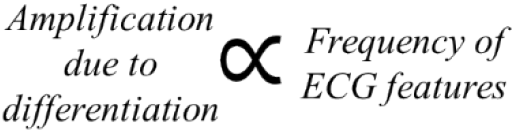 As R-waves have the highest frequency of the ECG features it is accentuated the most in the differentiated output (Fig 2A). The differentiation is applied via the following iteration,

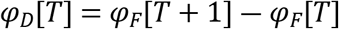

**Fig 2.**
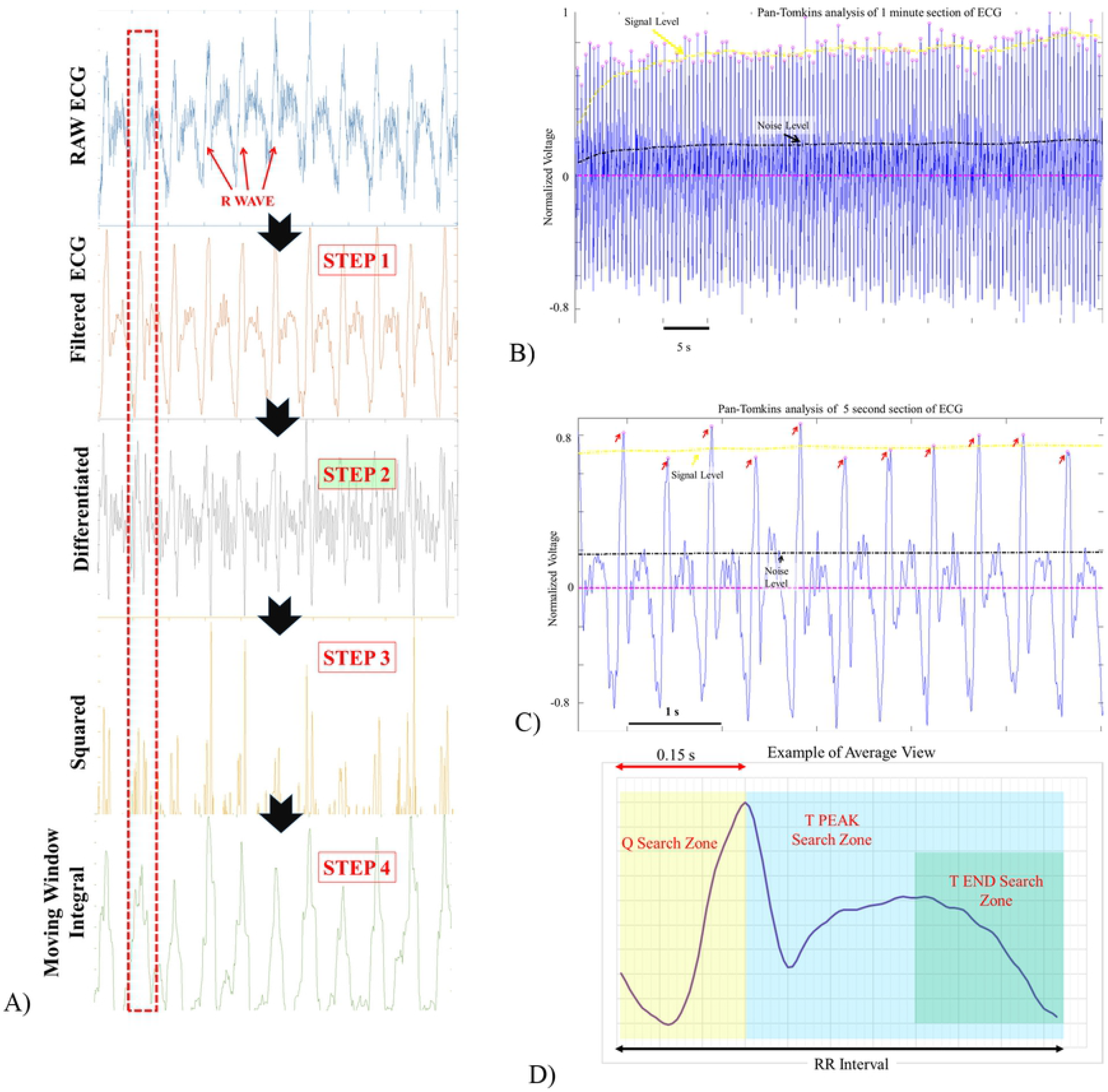
Graphical representations of the Pan-Tompkins algorithm applied the Larval ECG. A) The graphical outputs at each stage of the Pan-Tompkins algorithm, annotation are provided to show the position of the R-wave within the signal. B) The result of the Pan-Tompkins algorithm for a 1 minute section of data. Shown on the trace are the adaptive signal threshold and noise level utilized by the program to determine the position of each R-wave. C) A 5 second segment of ECG taken from the longer section to highlight the position of the fiducial points on each heartbeat. D) Example of the average waveform produced and the search zones that are used to find the position of the desired ECG characteristics Where *φ_D_* is the differentiated output and *φ_F_* is the filtered ECG signal. In MATLAB this can again be applied using an inbuilt function for which the code is given below,

%MATLAB ALGORITHM TO DIFFERENTIATE FILTERED ECG
%ecg_filt is the filtered data from the previous step
%ecg_diff is the differentiated output
ecg_diff=diff(ecg_filt);
3. In the thirds stage of the algorithm the differentiated output is squared by applying the iteration,

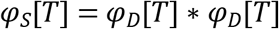 Where *φ_s_* is the squared ECG output. This step further accentuates the turning points in the ECG and makes the signal positive everywhere, as can be seen in Fig. 2A. As the P-T algorithm is specifically a QRS detection algorithm, the P and T-waves are usually relatively flat after this processing stage. The stage is performed using the following MATLAB code,

%MATLAB ALGORITHM TO SQUARE DIFFERENTIATED ECG
%ecg_diff is the differentiated data from the previous step
%ecg_sq is the squared output

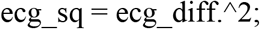
4. A moving window integral is then applied to the data that sums all the data points in a window of a defined width, Z. This is performed by applying the iteration;

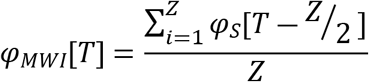 Where *φ_MWI_* is the integrated ECG output. If Z is roughly equal to the width of the QRS peak then it has the effect of creating large peaks in the region of the QRS, whilst ensuring that everything else is roughly zero. The output of this algorithm is shown in Fig. 2A.

%MATLAB ALGORITHM TO MOVING WINDOW AVERAGE SQUARED ECG
%ecg_sq is the squared signal from the previous step
%ecg_mwi is the moving window integral output

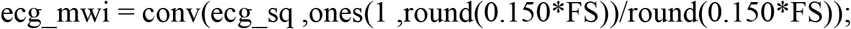
5. After the ECG has been processed in this way it is possible to detect the position of the QRS features via a peak-hunting algorithm. Pan-Tompkins developed a dual-threshold technique that adapts to the characteristics of the signal periodically to evaluate the ‘signal’ and ‘noise’ levels in the signal.

#### Pan-Tompkins Adaptive Thresholds

The algorithm first searches through the integrated waveform and an R wave is ‘detected’ every time a peak is found above the established threshold level. If the detected peak is below threshold, it is treated as a noise peak. If an R wave is not detected within 166% of the previously measured RR interval the program performs a search back using the second, lower threshold. Every time a peak is detected the algorithm determines if it is an R-wave using the previously established thresholds and updates the thresholds by the following algorithm:

If the peak that has been found is an R-wave then the new running estimate of the R peak height (R_PK_) is updated as,

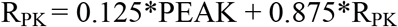

Where PEAK is the height of the detected peak.

If the peak that has been found is below threshold and therefore a noise peak, then the new running estimate of the noise peak heights (N_PK_) is updated as,

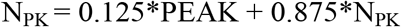

This enables the new thresholds are updated using;

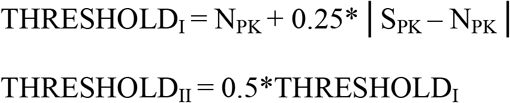

If a search back is used to find the peak then the new running estimate of the R-peak height is instead updated as,

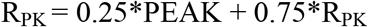

A sequence of values of R_PK_ and N_PK_ obtained for sections of data of different lengths are shown in Figs. 2B and 2C.

The program then searches through the filtered ECG signal to find QRS complexes using a similar approach. The program applies thresholds to the filtered ECG in the following way; if the detected peak (FPEAK) is an R-wave then the running estimate of the signal (FR_PK_) is updated as

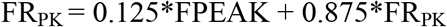

If the peak that has been found is a noise peak, then the new noise peak height (FN_PK_) is updated as,

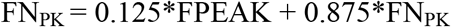

And again the new thresholds are updated as;

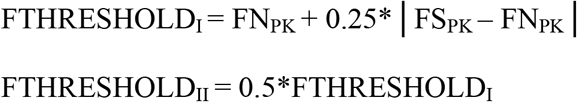

If a search back is used to find the peak then the new running estimate R-peak height is instead updated as,

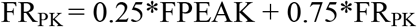

An identified QRS peak is only declared as an R-wave to be carried forward into the further analysis if it is detected in both passes. Every time a QRS peak is detected there is a 200 ms refractory period in which no other R-wave can be found.

#### Producing an Average Waveform from the Fiducial Points

Once the Pan-Tompkins algorithm has located the position of each R-wave in a section of ECG they can be used as fiducial points to produce an average waveform, 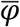. This is found by summing the voltage recorded at equivalent time points, T within each heartbeat, *i* and then dividing by the number of beats in the average. This enables the mean voltage at every position in the heart beat, relative to the R wave to be determined using the formula below,

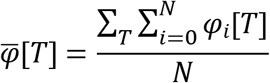

For the automated process the average waveform is calculated for a period that is equal to the measured RR interval that runs from 0.15 s before the R peak. This is shown in Fig. 2D.

### Detecting ECG features from the Average Waveform

To detect the features from the average waveform the program looks for peaks and troughs in appropriate windows within the heartbeat. The windows are shown in Fig. 2D and correspond to:

To find Q the software searches for the local minima in the region that is 0.15 s before the R peak. This window is guaranteed to contain a Q-wave if the R-wave frequency is greater than 6 Hz.

To find the T Peak the software searches for the local maxima that has the largest amplitude after the R peak. Once the T-peak has been found the program searches for the next point at which the ECG changes sign and designates this as the T end point. This is in accordance with other well established interpretations of the ECG signal [9].

### Calculating QTc from the Analyzed Waveform

The QT length is dependent on the heart rate of the fish and so it is necessary to calculate the corrected QTc by feeding in the measured QT and RR interval into the Bazett formula [10], where;

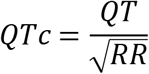

### Wavelet Transform Analysis

Wavelet transforms allow frequency analysis of a time-dependent signal, which allows the fundamental range of frequencies in the signal to be determined in order to optimize filter selection. The characteristics frequencies of the larval zebrafish ECG have never previously been shown and so it is desirable to evaluate them further. Applying Wavelet Transforms will thus aid the implementation of the Pan-Tomkins algorithm and aid future software designs.

A wavelet transform decomposes a signal into its fundamental parts that have a well-defined time and frequency localization. Through a convolution, the continuous wavelet transform (CWT) finds the inner product between the inputted signal and the analyzing wavelet that has a well-defined time-duration and frequency band. In a CWT the signal is compared to the analyzing wavelet that is time-shifted and scaled to yield coefficients that correspond to a measurement of the ECG constituents within the section and frequency band. In essence, the wavelet transform provides information about the ECG frequency at specific time-points within the heartbeat [11].

A CWT of a signal, *φ*(*t*) using a wavelet *μ*(*t*) is defined as,

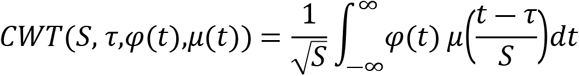

Where S is a scaling factor and τ is the position variable. The wavelet used to characterize the frequency of the ECG in this study is the ‘Mexican Hat’ wavelet [12].

To perform the CWT in MATLAB it is possible to utilize the inbuilt functions shown in the following code. The algorithm both performs the wavelet transform and plots the output together with original input signal.

%MATLAB ALGORITH TO PERFORM WAVELET TRANSFORM ON ECG

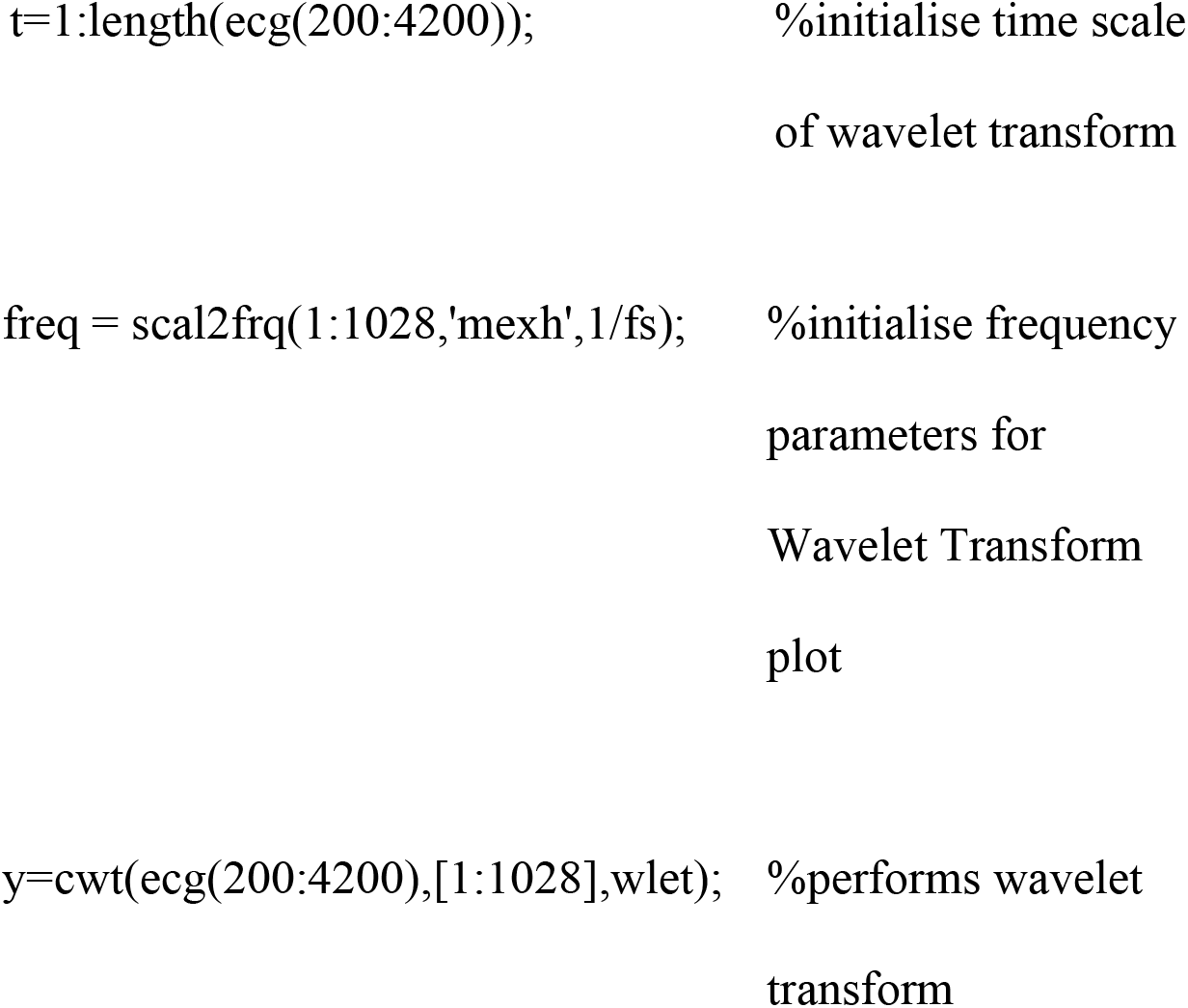
figure;
subplot(3,1,1);plot(t/fs,ecg(200:4200)),axis tight, title(‘Signal’); ylabel(‘Voltage’);
subplot(3,1,2:3);contour(t/fs,freq,abs(y)); axis tight, ylim([Filt_low,Filt_high]), xlabel(‘Time, s’),
ylabel(‘Frequency, Hz’),title(‘{\bf Wavelet Spectogram}’); colormap(‘default’);

### The frequency characteristics of the ECG

Contour plots of the Wavelet transform of the 20 second section of ECG and a shorter 1s section are shown in Fig. 3, together with the ECG trace on which the transform was performed. As can be seen in Fig. 3A, 42 heart beats were recorded in the section and from these beats the R-wave and T-wave frequency could be determined.

**Fig 3.**
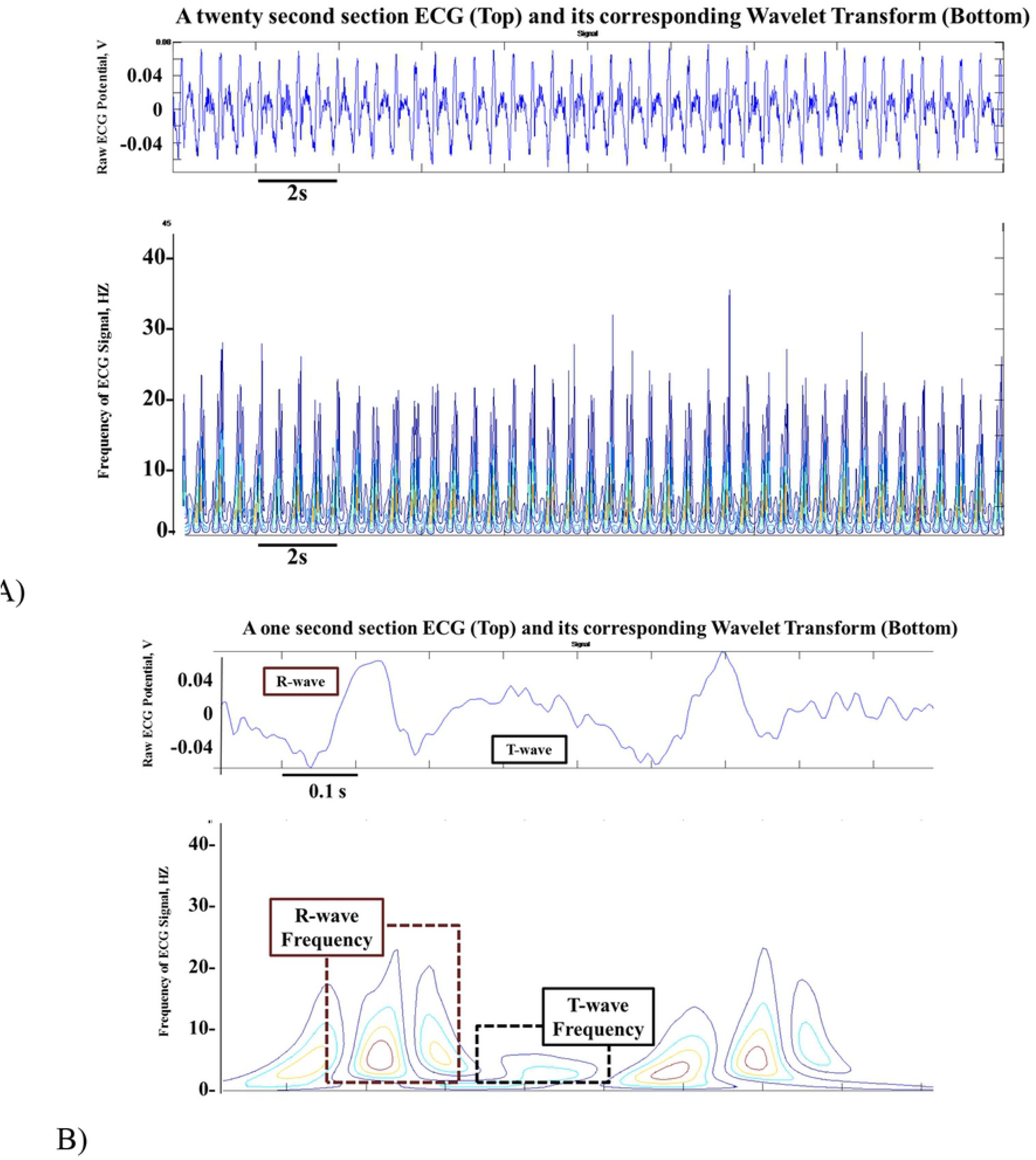
Contour plots to show the wavelet transforms of the ECG together with the Raw ECG trace. A) 20 seconds of normal zebrafish ECG, and its corresponding wavelet transform. B) 5 seconds of zebrafish ECG, and its corresponding wavelet transform. Annotations show the position of the R and T-waves and their corresponding frequencies

From the 42 beats the mean characteristic R-wave frequency was found to be, 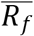 = 24.1 Hz, with a standard deviation of, σ = 3.4 Hz. The mean characteristic T-wave frequency was found to be, 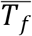 = 6.6 Hz, with a standard deviation of 1.3 Hz.

Assuming that most of the R-waves are adequately captured when the low pass filter is set to 2 standard deviations above mean frequency, a low pass cut-off of 31Hz was selected for the analysis. It can also be seen that a high pass filter of 1 Hz would not attenuate the signal. As a result of these observations it was decided that the data should be band pass filtered between 1 and 31 Hz to aid further analysis.

### LabChart^®^ Analysis

Fig. 4 shows a typical outcome of the LabChart^®^ analysis on a section of data. Immediately after the drug is added to the medium the measured QTc decreases which is followed by a steady increase in QTc until around 15 minutes after the drug has been administered. The maximum QTc change measured over this time is just under 5%. There is a steady increase in the RR interval from moment that the drug is administered from a minimum of 0.46 s to a maximum of around 0.7 s.

**Fig 4.**
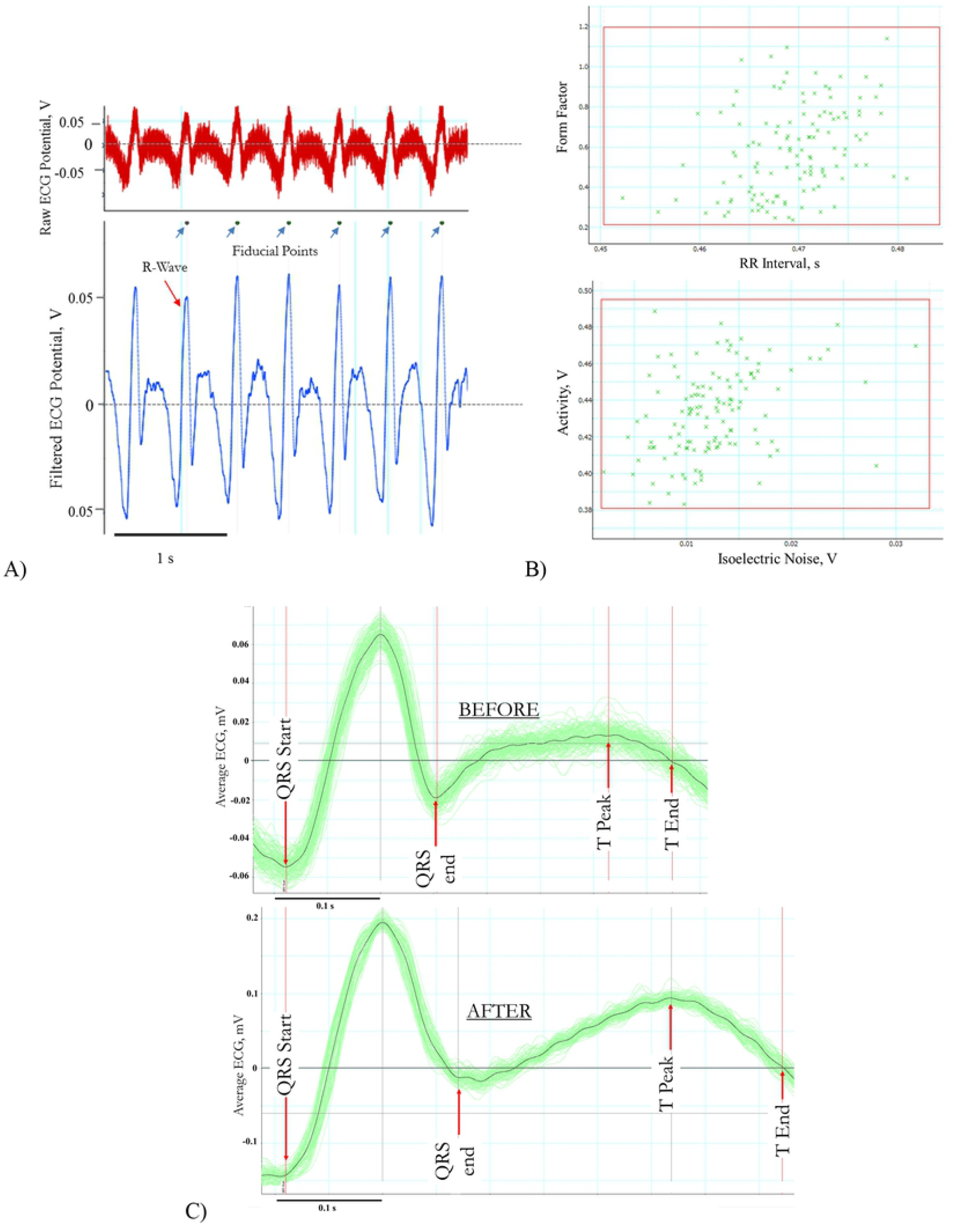
The LabChart^®^ analysis process of normal and pharmacologically altered larval ECG. The process consists of A) defining the position of each R-wave in the filtered ECG, B) Selecting which beats are to be ‘accepted’ to create C) the average ECG waveform. The average waveform generated for normal (before drug) and pharmacologically altered (25 minutes after drug delivery) ECG to highlight the change in the QT interval.

### The Automated Software Analysis

Our software was able to automatically detect the ECG peaks accurately, and drug effect on Q-T interval was determined (Fig. 5).

**Fig 5.**
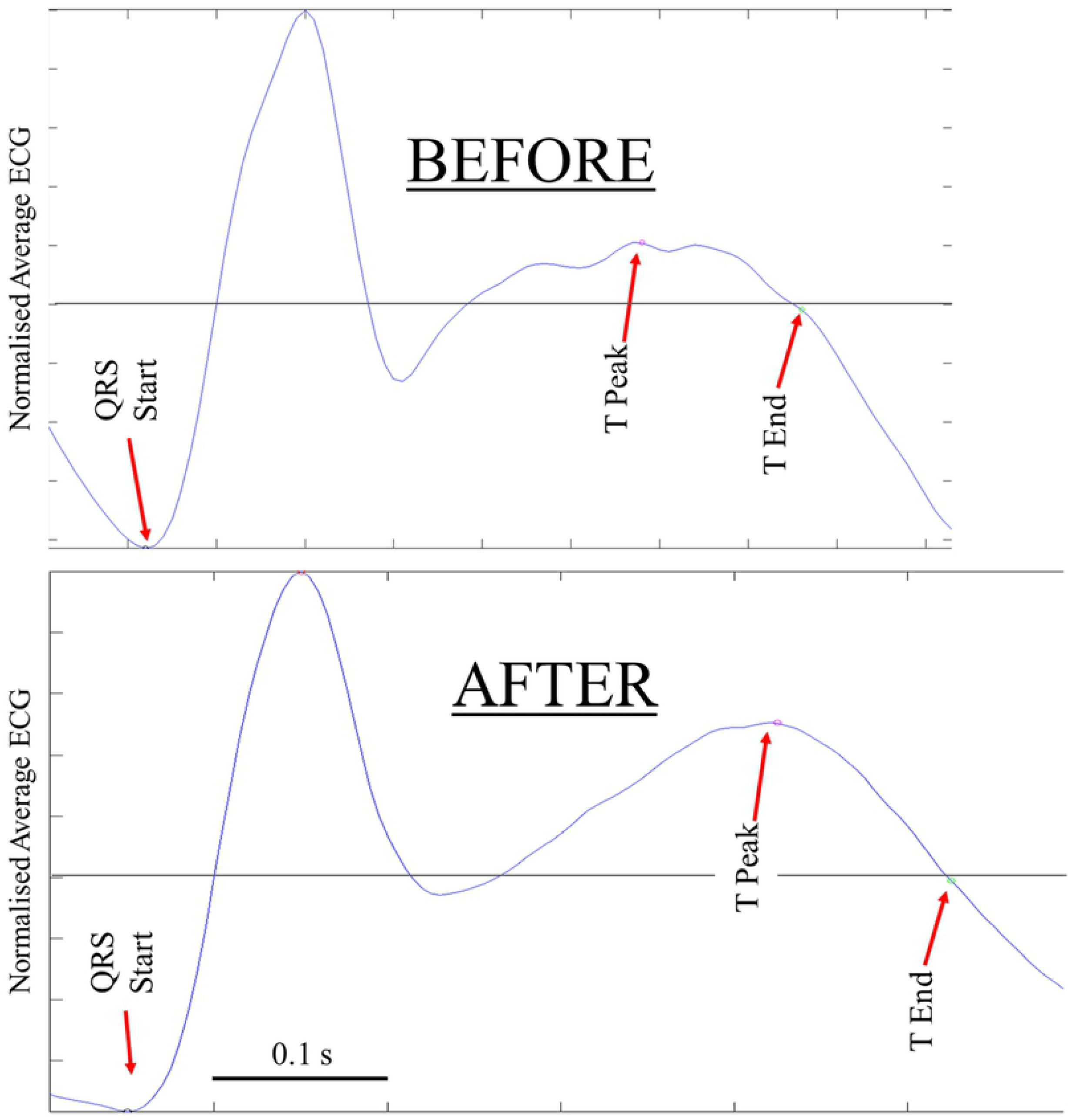
A comparison of the average ECG before and after 1mM of Verapamil had been added to the recording media. The top waveform produced from ECG 1 minute before the drug had been introduced, the bottom figure was taken from data 20 minutes after the drug had been introduced.

The measured QTc and RR interval from the automated analysis have been plotted in Fig. 6.

**Fig 6.**
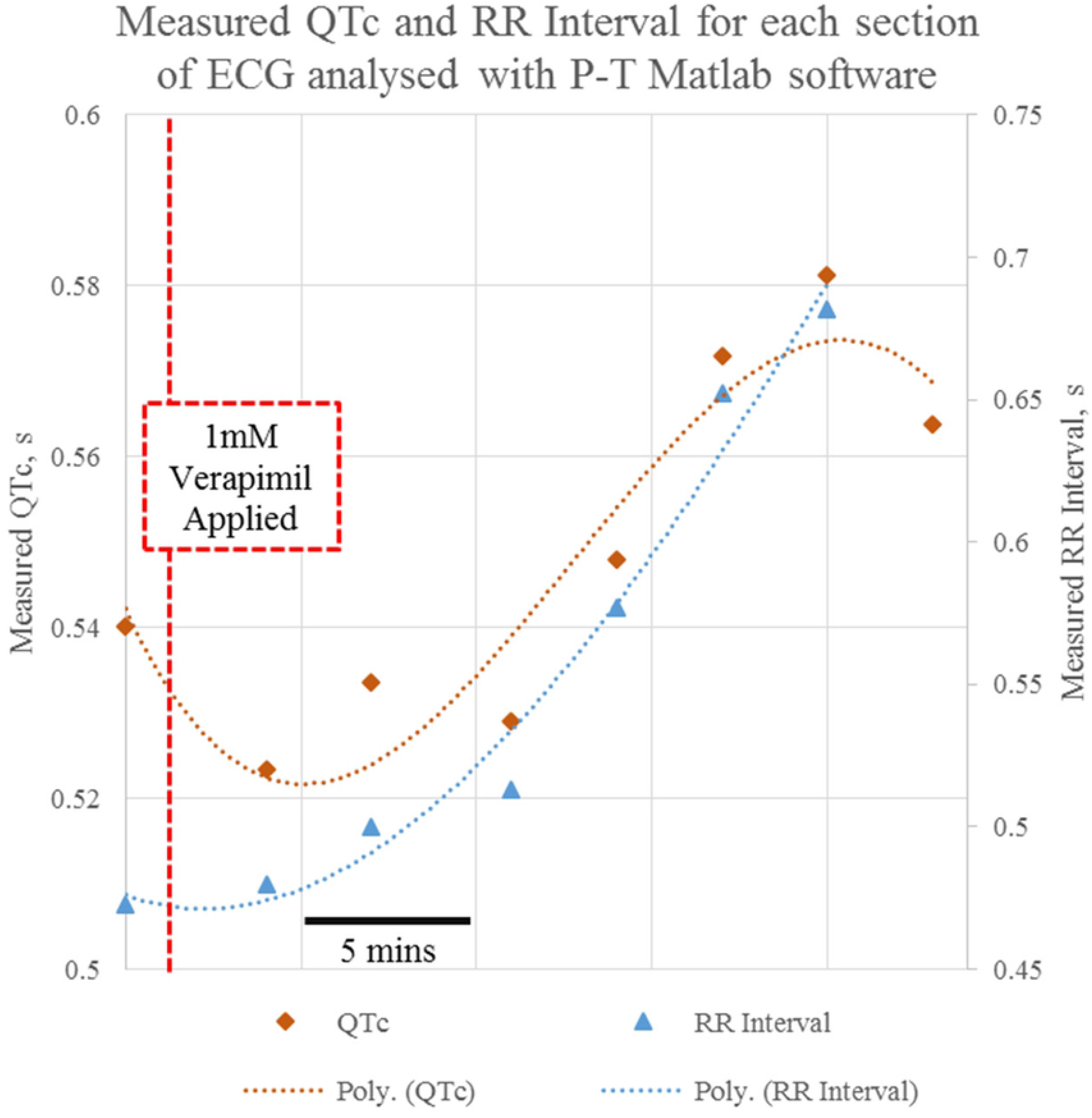
Scatter Graph to show the measured QTc and RR interval for the larvae from the MATLAB Software.

In the same way that was demonstrated by the LabChart^®^ analysis software, immediately after the drug is added to the medium the measured QTc decreases which is followed by an increase in QTc until around 15 minutes after the drug has been administered. However, for this analysis software the maximum QTc change measured over this time is just under 8%. It can also be seen that again there is a steady increase in the RR interval from moment that the drug is administered that is almost identical to the LabChart^®^ analysis software (Fig. 7).

**Fig 7.**
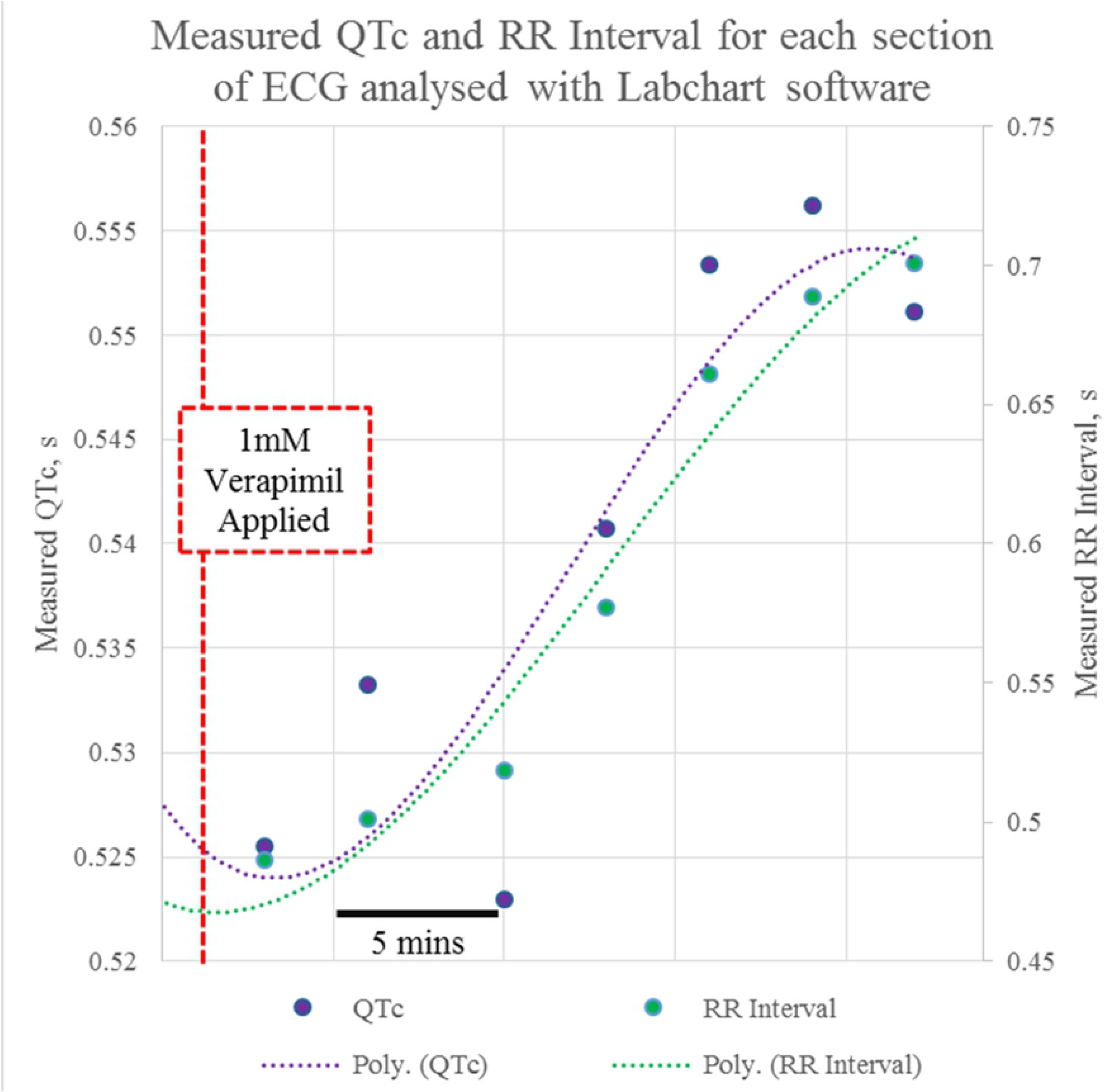
Scatter Graph to show the measured QTc and RR interval for the larvae from the LabChart^®^ Software.

### Comparison of the LabChart^®^ and MATLAB analysis software

To aid comparison between the two analysis techniques the LabChart^®^ and automated data are plotted together in Fig. 8. As can be seen in Fig. 8A, the measured QTc initially start off very similar until around 7 minutes after the drug has been added, after which the automated MATLAB software consistently records a higher QTc than the LabChart^®^ program. This phenomena occurs despite the programs measuring a very similar RR interval and detected number of beats for each section. The difference between the two analysis techniques is further illustrated in Fig. 8D, which shows the measured QTc change for both. Although the patterns are broadly similar, the automated process consistently records larger QTc change than the LabChart^®^ program.

**Fig 8.**
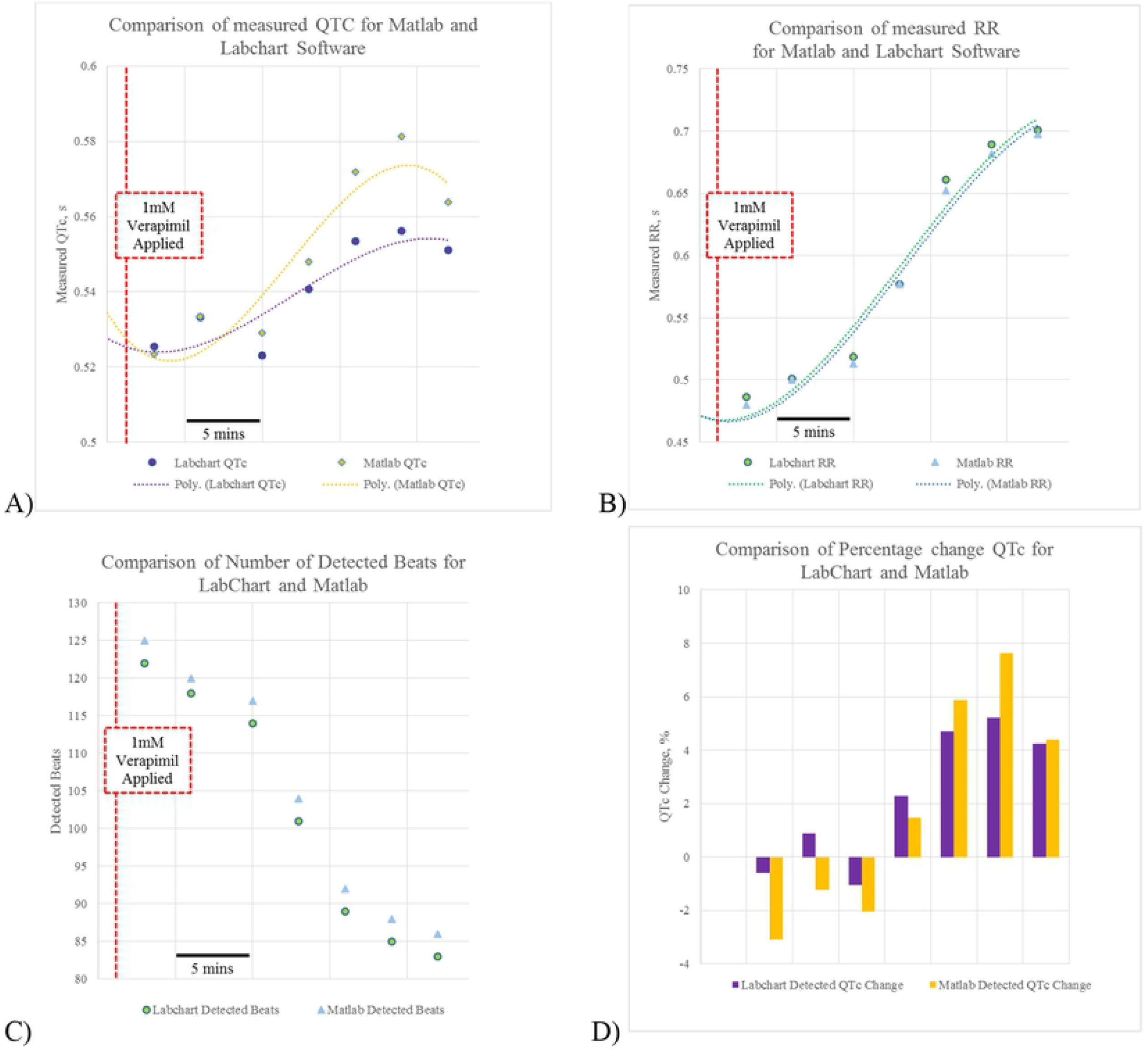
Plots to compare the measured ECG characteristics from LabChart^®^ and MATLAB software. A) Comparison of measure QTc B) Comparison of Measured RR interval C) Comparison of number of detected beats, D) comparison of measured QTc change for both programs.

## Discussion

The automated process is a significant improvement on the approaches that were previously applied to the zebrafish ECG. This article shows that the automated process that is based on established techniques of analyzing ECG can sensitively measure pharmacologically induced changes in the ECG.

However, it has also been shown that there are differences between the results obtained through the analysis with the LabChart^®^ software and the automated process. The main difference is that the automated process consistently measures a larger QTc change that the previously used approach. This can be explained by the fact the LabChart^®^ program analyses the whole section in one go, compared to the automated the process that outputs the QTc value for each 40 beats. Assuming that the QTc is not constant within each section of the ECG recording, the LabChart^®^ software will be less sensitive to subtle variations in the ECG as it instead finds a global average across the whole section. Even if QTc at the start and the end of the section are systematically different, the overall result will be somewhere in the middle. Instead, by focusing on smaller numbers of beats within the section the automated process provides a truer reflection of the actual QTc change within the recording and is more able to pick up these smaller changes.

The automated process is faster is able to detect the heart beats robustly and has less human involvement than the LabChart^®^ software, so it represents a significant improvement in the analysis available when analysis zebrafish larval ECG. Furthermore, there is no reason why this software cannot be applied to human ECG in the same way.

## Acknowledgments

This work was supported by the European Union’s Horizon 2020 Marie Sklodowska Curie Research and Innovation Staff Exchange programme (VISGEN, No. 734862) and OPEN FET RIA (NEURAM, No, 712821), the Higher Education Institutional Excellence Programme of the Ministry for Innovation and Technology in Hungary, within the framework of the “Innovation for the sustainable life and environment” thematic programme of the University of Pecs, and the Wellcome Trust Investigator Award (106955/Z/15/Z) to FM. The funders had no role in study design, data collection and analysis, decision to publish, or preparation of the manuscript.

